# Maternal cytokine response after SARS-CoV-2 infection during pregnancy

**DOI:** 10.1101/2022.01.04.474908

**Authors:** Frederieke A.J. Gigase, Nina M. Molenaar, Roy Missall, Anna-Sophie Rommel, Whitney Lieb, Erona Ibroci, Sophie Ohrn, Jezelle Lynch, Florian Krammer, Rachel I. Brody, Rebecca H. Jessel, Rhoda S. Sperling, Corina Lesseur, Francesco Callipari, Romeo R. Galang, Margaret C. Snead, Teresa Janevic, Joanne Stone, Elizabeth A. Howell, Jia Chen, Victor Pop, Siobhan M. Dolan, Veerle Bergink, Lotje D. De Witte

## Abstract

**Objective:** Dysregulation of the immune system during pregnancy is associated with adverse pregnancy outcomes. Recent studies report cytokine changes during the acute phase of severe acute respiratory syndrome coronavirus 2 (SARS-CoV-2) infection. We examine whether there is a lasting association between SARS-CoV-2 infection during pregnancy and peripheral blood cytokine levels.

**Study design:** We conducted a case-control study at the Mount Sinai health system in NYC including 100 SARS-CoV-2 IgG antibody positive people matched to 100 SARS-CoV-2 IgG antibody negative people on age, race/ethnicity, parity, and insurance status. Blood samples were collected at a median gestational age of 34 weeks. Levels of 14 cytokines were measured.

**Results:** Individual cytokine levels and cytokine cluster Eigenvalues did not differ significantly between groups, indicating no persisting maternal cytokine changes after SARS-CoV-2 infection during pregnancy.

**Conclusion:** Our findings suggest that the acute inflammatory response after SARS-CoV-2 infection may be restored to normal values during pregnancy.

## 1. Introduction

Coronavirus Disease 2019 (COVID-19), caused by the severe acute respiratory syndrome coronavirus 2 (SARS-CoV-2), continues to pose a global health threat. There is an urgent need to understand the impact of SARS-CoV-2 infection on populations at risk for severe COVID-19 illness, including pregnant and recently pregnant people. Pregnancy is characterized by complex changes in the immune system, making it a window of potential heightened susceptibility to various infections and increased risk of more severe illness once infected [1,2]. Recent studies have shown that SARS-CoV-2 infection during pregnancy is associated with increased risk for severe infection and adverse pregnancy outcomes, including preterm birth (PTB), low birth weight, and pre-eclampsia [3,4,5,6]. Another recent study did not find an effect of past maternal SARS-CoV-2 infection on obstetric and neonatal outcomes [7].

Accumulating evidence suggests that excessive cytokine production contributes to disease severity and adverse outcomes in patients with COVID-19 [8]. In cases with severe SARS-CoV-2 infection, high levels of inflammatory markers such as C-reactive protein (CRP), cytokines and chemokines such as interleukin (IL)-2, IL-6, IL-10, and tumor necrosis factor α (TNFα), and a high neutrophil to lymphocyte ratio have been observed [9,10,11]. In addition to an acute immune response, immune changes may persist after SARS-CoV-2 infection. Files et al. showed that immune changes of B and T cell phenotypes persisted at 30 days after confirmed SARS-CoV-2 infection in patients who had since recovered, suggesting a prolonged period of immune dysregulation after SARS-CoV-2 infection of which the consequences are not yet clear [12]. Immune changes play a regulatory role during pregnancy [1]. In early pregnancy, these changes contribute to successful implantation and decidualization [13]. In later pregnancy, inflammatory pathways are involved in myometrial quiescence and activation, cervical ripening, and weakening of the fetal membrane [14]. Dysregulation of the immune system during pregnancy has been associated with adverse pregnancy outcomes, including miscarriage, intrauterine growth restriction (IUGR), and PTB, and can be detrimental for long-term neonatal outcomes [15].

Three recent studies have analyzed the impact of acute SARS-CoV-2 infection on the maternal immune system. Tanacan et al. showed that acute SARS-CoV-2 infection during pregnancy was associated with increased peripheral blood levels of interferon (IFN)-γ in the third trimester and decreased levels of IL-2 in the first and second trimester, and IL-10 and IL-17 in the first trimester (N=180) [16]. Another retrospective study showed that plasma cytokine levels of IL-2, IL-5, IL-10 and IL-12 were similar in pregnant and non-pregnant SARS-CoV-2 infected people, and were decreased compared to healthy pregnant people (N=25) [17]. Sherer *et al*. found that gene expression of *IL-1β* and *IL-6* in whole blood samples of pregnant people at delivery was not altered within 14 days after a positive SARS-CoV-2 test compared to pregnant people with a negative test (N=33) [18]. Although previous studies suggest peripheral blood cytokine changes in response to acute SARS-CoV-2 infection during pregnancy, they do not fully align with each other, which may be due to limited sample size, varying study design or other factors influencing cytokine levels including nutrition, BMI, and previous infections. Further studies are needed to clarify these results and to understand whether SARS-CoV-2 infection during pregnancy is associated with persisting immune changes.

In the current study, we investigated whether SARS-CoV-2 IgG antibody status during pregnancy is associated with lasting immune changes to better understand the interaction between infection and inflammation during pregnancy. A prospective pregnancy cohort was established at the Mount Sinai Health System (MSHS) in New York City (NYC) at the beginning of the pandemic in April 2020. The current interim analysis was performed on 200 participants as the COVID-19 pandemic is ongoing with an urgent need for data to understand the effect of SARS-CoV-2 infection during pregnancy. We measured IgG antibodies to the SARS-CoV-2 spike (S) protein to analyze past exposure to SARS-CoV-2 infection [19,20]. IgG antibodies to the S protein are detectable between 7 and 30 days after infection and maintained for at least three months after infection [21,22]. The plasma levels of 14 cytokines were measured in 100 IgG antibody positive and 100 IgG antibody negative participants. As cytokines are not regulated independently of each other [23], we compared groups for both the level of individual cytokines and cytokine clusters.

## 2. Material and Methods

### 2.1 Study design and participants

The Generation C Study is a prospective pregnancy cohort study that is being conducted at the MSHS, the largest healthcare system in NYC. The Generation C study is designed to examine the impact of SARS-CoV-2 infection during pregnancy on obstetric and neonatal outcomes and the mediating role of the immune response. All pregnant people receiving obstetrical care at the Mount Sinai Hospital and Mount Sinai West during the study period are eligible for participation. Recruitment and sampling started on April 20, 2020 and is currently ongoing. Exclusion criteria were age under 18 years. Participants enrolled in the Generation C study are seen multiple times during pregnancy and blood is collected as part of routine clinical care. Serological assays are used to confirm past SARS-CoV-2 infection.

For this interim analysis, we conducted a case-control study of people with and without evidence of recent SARS-CoV-2 infection based on detection of IgG antibodies against the SARS-CoV-2 spike (S) protein (anti-S IgG). The cases consisted of 100 pregnant people with anti-S IgG antibodies. They were matched with a control group of 100 pregnant people with no detectable anti-S IgG antibodies, on age, race/ethnicity, parity, and insurance status. All participants in this study gave birth to a singleton infant between April 24 and September 28, 2020. The first COVID-19 case in NYC was officially confirmed on March 1, 2020, and seropositive cases peaked in the metropolitan area in March and April 2020 [24]. Therefore, it was inferred that the cases in the current study were most likely exposed to SARS-CoV-2 during pregnancy, prior to the first COVID-19 vaccine receiving Emergency Use Authorization in the U.S. on December 11, 2020. Blood samples were obtained as part of the routine blood draw during the prenatal visits or admisstion to labor and delivery. The latest available blood sample obtained at routine blood draw or at labor and delivery was used for analysis. Relatively more third trimester samples were available due to the timing of the study and the analysis of the latest available blood sample – which was in the third trimester for most participants. Participants were not given dietary restrictions prior to blood draw. Blood was extracted into 4 ml lavender top ethylene damine tetra acetic acid (EDTA) vacutainer tubes. Samples were centrifuged at 1300 rpm (350xg) for 10 minutes at room temperature. Plasma aliquots were stored in cryogenic tubes at -80 until further analysis. The blood samples were collected either in the second (cases: n=7, controls: n=8) or third trimester (cases: n= 56, controls: n= 54), or upon admission to labor and delivery (cases: n=37, controls: n=38), at a median gestational age of 36 weeks (cases: 36 weeks, SD 4.9, controls: 36 weeks, SD 5.8 weeks). These samples were used for two purposes: 1) to establish anti-S IgG antibody titer level and 2) for multiplex cytokine analysis. All participants provided written informed consent per the institutional review board (IRB)-approved study protocol (IRB at the Icahn School of Medicine at Mount Sinai, protocol IRB-20-03352, April 15, 2020). This activity was reviewed by CDC and was conducted consistent with applicable federal law and CDC policy.

### 2.2 Serological testing

Serological testing was performed using a serologic enzyme-linked immunosorbent assay (ELISA) developed at the Icahn School of Medicine at Mount Sinai [20]. The assay has a relatively high sensitivity (95%) and specificity (100%) and a positive predictive value of 100%, with a negative predictive value of 97% [20], minimizing misclassification. The plasma samples were first screened using a low dilution (1:50) for antibodies to SARS-CoV-2 receptor binding domain (RBD). If plasma samples tested positive for RBD, they were further diluted and tested in an assay using the full-length spike protein to validate positivity (titer > 0) and to determine the anti-S IgG endpoint titer. A general limitation in SARS-CoV-2 IgG testing is the possibility of false negatives prior to IgG antibody production onset or in cases where IgG antibody levels revert to zero in the months after infection [25].

### 2.3 Multiplex analysis of cytokines

The same plasma sample was used to quantify the levels of 14 cytokines, through the High Sensitivity T-Cell Discovery Array 14-Plex (Millipore, St. Charles, MO, USA) according to the manufacturer’s protocol. The multiplex assay was performed at Eve Technologies using the Bio-Plex™ 200 system (Bio-Rad Laboratories, Inc., Hercules, CA, USA). The 14-plex consisted of granulocyte-macrophage colony-stimulating factor (GM-CSF), IFN-γ, IL-1β, IL-2, IL-4, IL-5, IL-6, IL-8, IL-10, IL-12 (p70), IL-13, IL-17A, IL-23, and TNFα. This panel was selected because it contains cytokines that have been identified as part of the cytokine storm in response to severe SARS-CoV-2 infection [9], as well as cytokines that have been related to adverse pregnancy outcomes [26,27,28]. The assay sensitivities of these markers range from 0.11 – 3.25 pg/mL. Individual analyte values and other assay details are available on Eve Technologies’ website or in the Milliplex protocol. The performance of the assays is shown in Supplementary Table I.

### 2.4 Demographic and clinical characteristics

Demographic and clinical characteristics were extracted from electronic medical records (EMR). A 50g oral glucose tolerance test (GTT) is performed as part of routine prenatal care in the MSHS. If > 140mg/dl a confirmatory 3 hour GTT is performed and if > 2 values are abnormal, a woman is diagnosed with gestational diabetes. Analyses were adjusted for covariates that are potential risk factors for either SARS-CoV-2 infection and/or cytokine changes using step-wise procedures, including: age, pre-pregnancy body mass index (BMI), race/ethnicity, parity, gestational age, gestational diabetes and time between blood draw and processing [29,30,31,32,33].

### 2.5 Statistical analysis

Demographic and clinical characteristics of cases (100 pregnant people with anti-S IgG antibodies) and controls (100 pregnant people with no anti-S IgG antibodies) were compared using either an independent samples t-test or a Mann-Whitney U test for continuous variables and a chi-square test for categorical variables. Correlations were assessed using Spearman’s rho. Cytokines with negative or non-significant correlations with other cytokines were excluded from further analysis. Prior to clustering, each cytokine was scaled and centered; clusters were identified using complete-linkage agglomerative hierarchical clustering with Euclidean distances across cases and controls using the pheatmap v1.0.2 package in R [34]. The optimal number of clusters was confirmed using the elbow-method by calculating cluster-sum of squared errors (wss) for multiple values of k with the fviz_nclust function from the factoextra package in R [35]. Bartlett’s test of sphericity and the Kaiser-Meyer-Olkin (KMO) measure of sampling adequacy (MSA) were used to test suitability of the implementation of a principal component analysis on the data [36]. The factor loading cut-off was set at an absolute value of 0.5. For each cytokine cluster, we performed principal component analysis, the first principal component or cluster Eigenvalue summarizes the cluster-specific cytokine expression across samples [37]. The association of the cytokines with their respective cluster Eigenvalue was assessed using correlations. Plots were generated in R using the pheatmap, ggplot2 and corrplot packages, and in GraphPad Prism.

Multiple linear regression analyses were conducted to examine the association of anti-S IgG status with individual cytokine levels and cluster Eigenvalues. Multiple linear regression models were adjusted in a step-wise procedure for covariates described above. To control for the cytokine changes that may occur during active labor, a sensitivity analysis was performed removing participants (both cases and controls) with samples taken at labor and delivery from the analysis. In addition, linear regression analyses were performed separately for samples from the 3^rd^ trimester and labor and delivery samples. Bonferroni correction was applied to correct for the effect of multiple testing (p<.007). Antibody titer level has been associated with disease severity and was used as a proxy for disease severity [38]. To examine the association between cytokine levels and antibody titer levels, cases were divided into three groups based on anti-S IgG antibody titer level for ease of interpretation: mild (1:80-1:400), moderate (1:800-1:1600), and high (>1:1600). Individual cytokine levels and cluster Eigenvalues were compared across titer groups using a Kruskall Wallis test. Statistical analyses were performed using SPSS software version 27.0.

## 3. Results

### 3.1 Clinical characteristics

Participants in this case-control study delivered at a median gestational age of 39 weeks (cases: 39.1 weeks, SD 14 days, controls: 39 weeks, SD 12 days). The median gestational age at blood draw was 36 weeks (cases: 36 weeks, SD 5 weeks, controls: 36 weeks, SD 5.8 weeks). The average number of days between blood draw and delivery was 30 days (cases: 28 days, SD 32 days, controls: 32 days, SD 37 days). Demographics and clinical characteristics were not significantly different between cases and controls (Table I). Among cases, antibody levels were mild, moderate, and high in 37 (37%), 47 (47%), and 16 (16%) participants, respectively.

**Table I.** Comparison of characteristics of 200 pregnant people by SARS-CoV-2 status. A case-control study was performed at the Mount Sinai Health System in NYC. Demographic characteristics were compared between cases (100 pregnant people with anti-S IgG antibodies) and controls (100 pregnant people with no anti-S IgG antibodies). Chi-squared tests were performed for race/ethnicity, insurance, parity, smoking, hypertension, diabetes and gestational diabetes. Demographic characteristics did not significantly differ between cases and controls.

| Demographic Characteristics | Controls (N=100) | Cases (N=100) | *p*-value |
| --- | --- | --- | --- |
| Maternal age in years, mean (SD)<sup>a</sup> | 33 (5) | 33 (5) | .62 |
| Race/ethnicity, n<sup>b</sup> |  |  | .11 |
| Asian, non-Hispanic | 7 | 2 |  |
| Black/African American, non-Hispanic | 21 | 25 |  |
| Hispanic | 32 | 46 |  |
| White, non-Hispanic | 35 | 22 |  |
| Other, non-Hispanic | 3 | 4 |  |
| Unknown | 2 | 1 |  |
| Insurance, n<sup>c</sup> |  |  | .28 |
| Private | 61 | 50 |  |
| Public | 27 | 39 |  |
| Self-pay | 1 | 2 |  |
| Other/unknown | 11 | 9 |  |
| Parity, n<sup>d</sup> |  |  | .57 |
| Nulliparous | 44 | 40 |  |
| Multiparous | 56 | 60 |  |
| Pre-pregnancy body mass index (kg/m<sup>2</sup>), | 27.6 (7.3) | 28.4 (5.4) | .53 |
| mean (SD)<sup>a</sup> |  |  |  |
| Smoking, n<sup>e</sup> |  |  | .54 |
| Smoking during pregnancy | 0 | 0 |  |
| Smoking prior to pregnancy | 10 | 8 |  |
| Non smoker | 90 | 92 |  |
| Pre-pregnancy hypertension, n<sup>f</sup> | 3 | 2 | .65 |
| Pre-pregnancy diabetes, n<sup>g</sup> | 1 | 1 | 1.0 |
| Gestational diabetes, n<sup>h</sup> | 5 | 6 | .46 |

<sup>a</sup>: Independent samples t-test
<sup>b</sup>: Chi-squared test; DF = 5
<sup>c</sup>: Chi-squared test; DF = 3
<sup>d</sup>: Chi-squared test; DF = 1
<sup>e</sup>: Chi-squared test; DF = 1
<sup>f</sup>: Chi-squared test; DF = 2
<sup>g</sup>: Chi-squared test; DF = 2
<sup>h</sup>: Chi-squared test; DF = 2

### 3.2 Association between anti-S IgG status and individual cytokine levels

Cytokine values for IL-4 and IL-13 were not detected in two samples (one case and one control sample) and these were excluded from the analysis (Supplementary Table I). All other plasma samples (n=198) showed cytokine values detected within the standard curve. The distribution of cytokine levels were compared between cases and controls (Figure 1); no significant differences were noted. We found no association between anti-S IgG status and cytokine levels, in either unadjusted or adjusted analyses (Table II). In addition, we found no relationship between anti-S IgG antibody titer levels and cytokine levels among cases (Supplementary table II). To control for the effect of active labor on cytokine levels, a sensitivity analysis (n=125) was performed excluding samples obtained at labor and delivery (n=75), which showed similar results for cytokine levels compared between cases and controls (Supplementary table III). To assess the effect of gestational age on cytokine levels, linear regression analyses were performed separately for samples from the 3^rd^ trimester and at labor and delivery, which showed no significant difference in cytokine levels between cases and controls in the 3rd trimester, and significantly lower IL-23 in cases compared to controls at labor and delivery (Supplementary Table IV and V).

**Figure 1.**
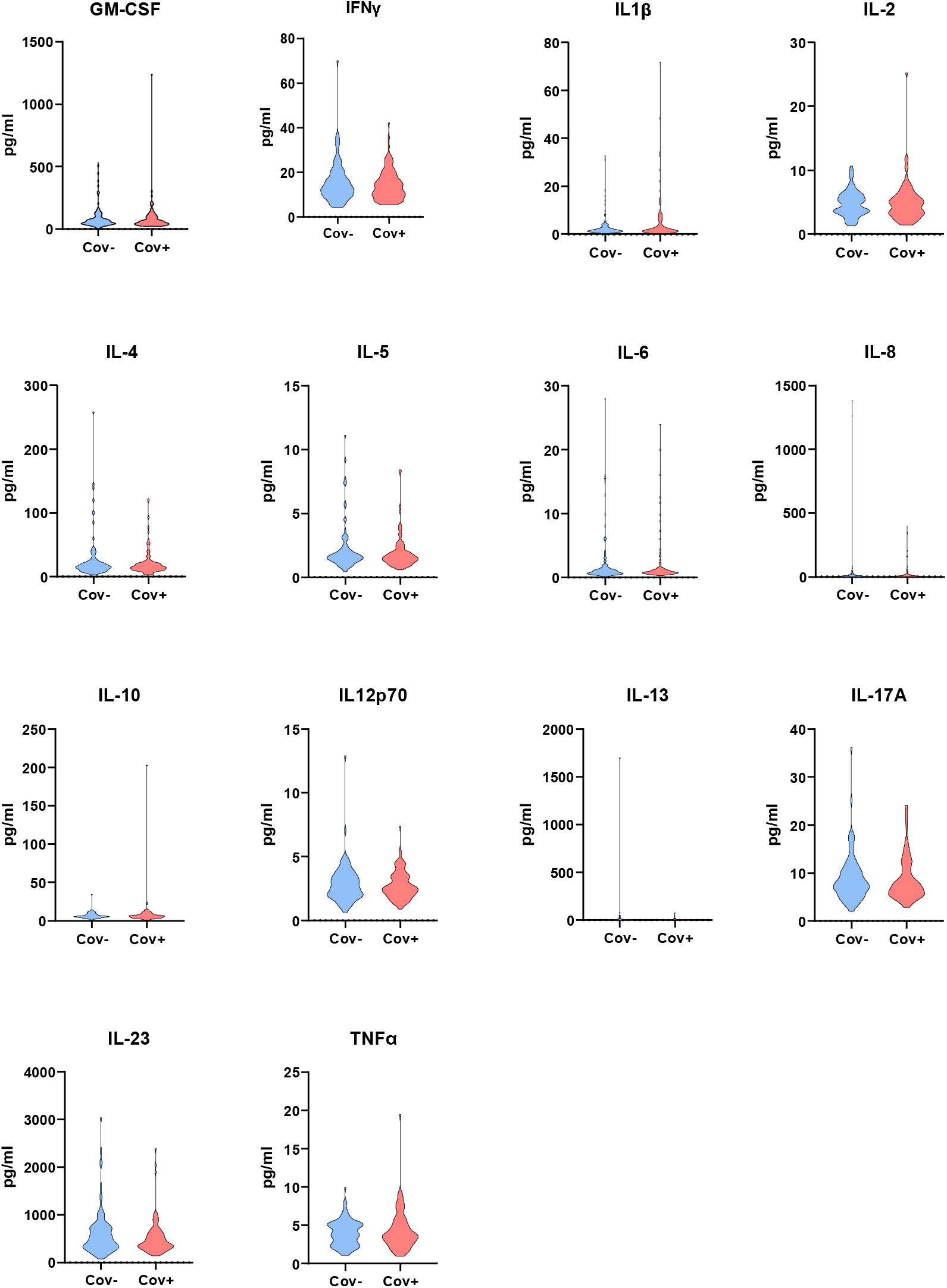
Peripheral blood levels (pg/ml) of 14 cytokines in cases (pregnant people with anti-S IgG antibodies, n=99) and controls (pregnant people with no anti-S IgG antibodies, n=99) referred to as cov+ and cov-respectively. Distribution of the cytokine levels are comparable between cases and controls. Pg = picogram, GM-CSF = granulocyte-macrophage colony-stimulating factor, IFN-γ = Interferon, IL = interleukin, TNF = tumor necrosis factor.

**Table II.** Association between peripheral blood cytokine levels and SARS-CoV-2 status among pregnant people. Peripheral blood cytokine levels were compared between cases (pregnant people with anti-S IgG antibodies, n=99) and controls (pregnant people with no anti-S IgG antibodies, n=99). Linear regression analyses were performed with plasma cytokine levels (pg/ml) as the dependent variables and SARS-CoV-2 status (anti-S IgG antibody positive / negative) as the exposure variable. Analyses were adjusted for covariates listed below. Cytokine levels were not significantly different between cases and controls. GM-CSF = granulocyte-macrophage colony-stimulating factor, IFN-γ = Interferon, IL = interleukin.

| Cytokine | Unadjusted $\beta$ | 95% CI | *p*-value | Adjusted $\beta^a$ | 95% CI | *p*-value |
| --- | --- | --- | --- | --- | --- | --- |
| GM-CSF | -.02 | -.11; .07 | .63 | .00 | -.09; .09 | .99 |
| IFN-γ | -.02 | -.08; .04 | .57 | .009 | -.05; .07 | .76 |
| IL-1β | .08 | -.04; .2 | .18 | .06 | -.04; .17 | .22 |
| IL-2 | -.01 | -.07; .05 | .82 | .03 | -.03; .08 | .4 |
| IL-4 | -.02 | -.11; .07 | .69 | .003 | -.09; .1 | .95 |
| IL-5 | -.03 | -.1; .03 | .32 | -.03 | -.09; .05 | .48 |
| IL-6 | -.01 | -.13; .1 | .84 | -.02 | -.14; .1 | .76 |
| IL-8 | .05 | -.09; .19 | .52 | -.03 | -.15; .1 | .7 |
| IL-10 | .01 | -.06; .08 | .74 | .03 | -.04; .11 | .42 |
| IL-12p70 | .003 | -.05; .06 | .91 | .02 | -.03; .07 | .46 |
| IL-13 | -.07 | -.19; .06 | .31 | -.06 | -.2; .07 | .36 |
| IL-17A | -.01 | -.07; .05 | .79 | .02 | -.04; .08 | .53 |
| IL-23 | -.05 | -.13; .02 | .15 | -.04 | -.11; .04 | .34 |
| TNFα | .02 | -.04; .08 | .52 | .01 | -.05; .07 | .82 |

<sup>a</sup>: Adjusted for: maternal age, race/ethnicity, pre-pregnancy body mass index, gestational diabetes, parity, gestational age in weeks at blood draw, being in labor at the time of blood draw (yes / no), and time between blood draw & sample processing (hours).

### 3.3 Association between anti-S IgG status and cluster Eigenvalues

Spearman correlation analyses showed positive correlations (range rho = 0.15-0.8) between most of the cytokines, with the exception of TNFα (rho ranging between -0.29 and 0.39) (Figure 2; Supplementary table VI). TNFα was expected to show a positive correlation with other cytokines. We assessed whether biological mechanisms interfered with TNFα levels, but this was not the case for time between blood draw and processing (r=.095, p=.18) or being in labor at the time of blood draw (r=-.07, p=.35). Hierarchical cluster analysis was performed including 13 correlated cytokines. Three clusters were identified: cytokine cluster 1 (IL-1β, IL-8) (r^2^ = 11.4%, MSA = .5, Bartlett’s p<.001); cytokine cluster 2 (IL-4, IL-5, IL-6, IL-13) (r^2^ = 13.7%, MSA = .832, Bartlett’s p<.001); and cytokine cluster 3 (IL-10, IL-12p70, IL-17A, IL-23, IFN-γ, IL-2, GM-CSF) (r^2^ = 30.7%, MSA = .879, Bartlett’s p<.001), with a cumulative 55.77% variance explained (Supplementary table VII). The cytokines in cluster 1, 2 and 3 show the strongest correlation with their respective cluster Eigenvalue compared to the individual cytokines, indicating that the cluster Eigenvalues sufficiently capture the activity of each cytokine within that cluster (Supplementary Figure 1; Supplementary Figure 2). We next compared the cluster Eigenvalues between cases and controls. Hierarchical clustering revealed no association between anti-S IgG status and cytokine clustering (Figure 3). Indeed, we found no association between anti-S IgG status and cluster Eigenvalues, both in unadjusted and adjusted analyses (Table III). In addition, we found no relationship between anti-S IgG antibody titer levels and cluster Eigenvalues among cases (Supplementary table VIII). The sensitivity analysis excluding participants in active labor showed similar results for cluster Eigenvalues compared between cases and controls (Supplementary table IX).

**Figure 2.**
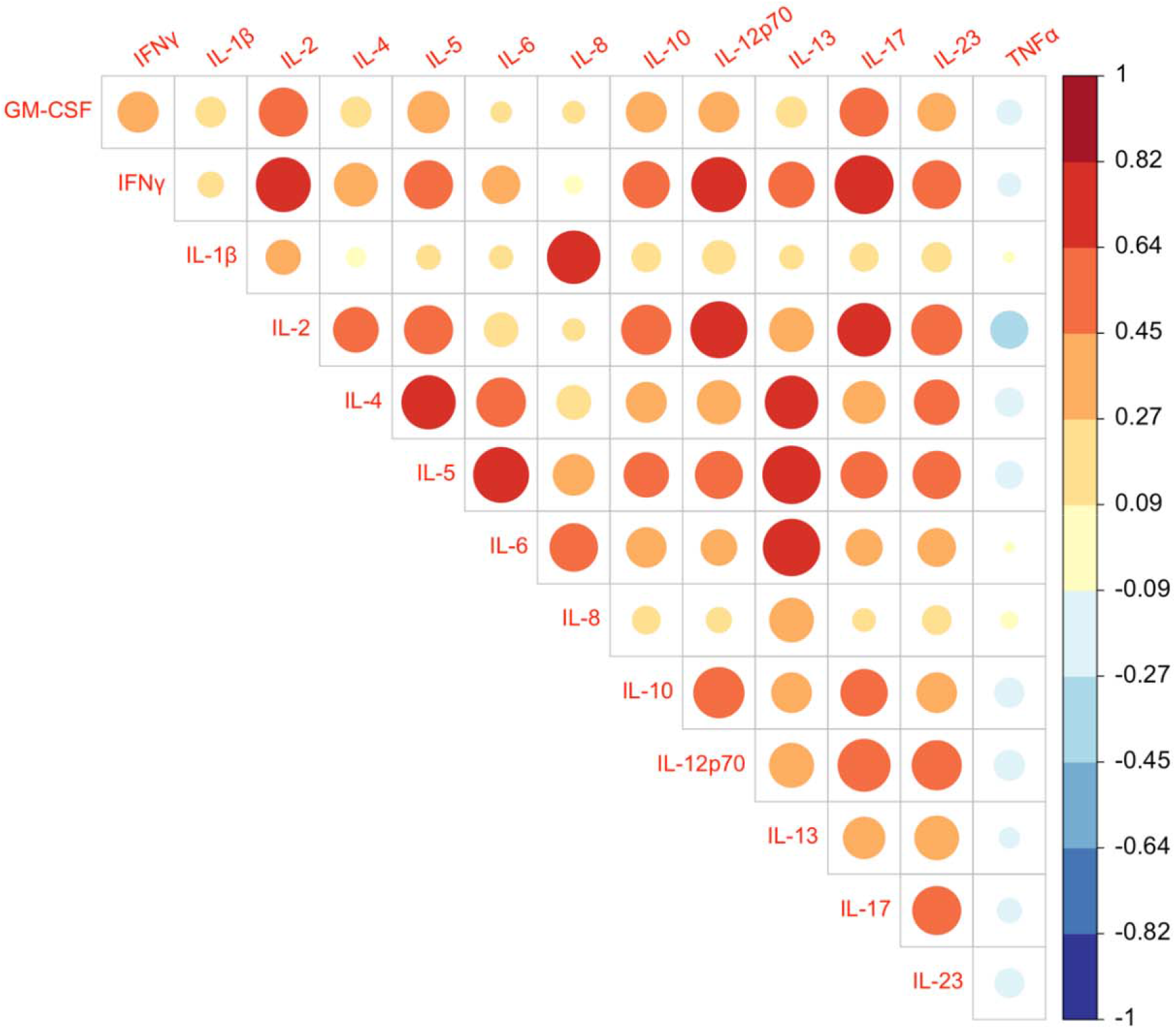
Correlation plot of 14 cytokines assessed in the peripheral blood of cases (pregnant people with anti-S IgG antibodies, n=99) and controls (pregnant people with no anti-S IgG antibodies, n=99). Darker colors indicate stronger positive (red) or negative (blue) Spearman correlations. Almost all correlations were significant (p<.005). TNFα shows a non-significant and mostly negative correlation with other cytokines (rho ranging between -0.288 and 0.39). The p-values of the correlations are shown in Supplementary Table VI. GM-CSF = granulocyte-macrophage colony-stimulating factor, IFN-γ = Interferon, IL = interleukin, TNF = tumor necrosis factor.

**Figure 3.**
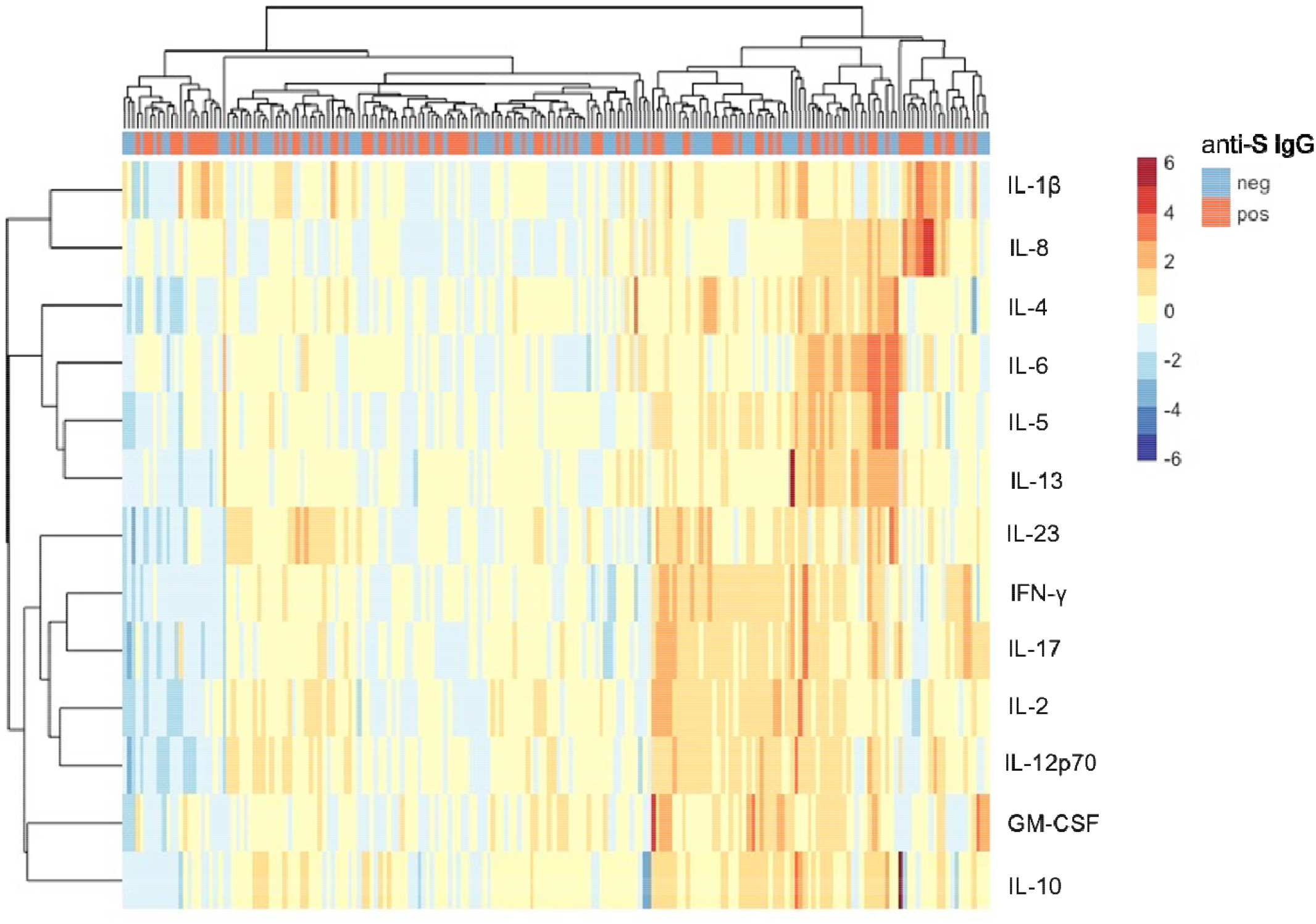
Heatmap showing hierarchical clustering analysis of 13 correlated cytokines measured in cases (pregnant people with anti-S IgG antibodies, n=99) and controls (pregnant people with no anti-S IgG antibodies, n=99). Scale are row-scaled (z-standardized) cytokine expression values. Darker colors indicate higher (red) or lower (blue) expression levels than the respective mean. Distance measures show no sample clustering (column) based on anti-S IgG antibody status. Three cytokine clusters (rows) can be identified. GM-CSF = granulocyte-macrophage colony-stimulating factor, IFN-γ = Interferon, IL = interleukin, neg = negative (controls), pos = positive (cases).

**Table III.** Association between peripheral blood cytokine clusters and SARS-CoV-2 status among pregnant people. Eigenvalues of three cytokine clusters were compared between cases (pregnant people with anti-S IgG antibodies, n=99) and controls (pregnant people with no anti-S IgG antibodies, n=99). Linear regression analyses were performed with the Eigenvalues of three cytokine clusters as the dependent variables and SARS-CoV-2 status (anti-S IgG antibody positive / negative) as the exposure variable. Analyses were adjusted for covariates listed below. Cytokine clusters were not significantly different between cases and controls. GM-CSF = granulocyte-macrophage colony-stimulating factor, IFN-γ = Interferon, IL = interleukin.

| Cluster | Cytokines in cluster | Unadjusted<br>$\beta$ | 95% CI | *p*-value | Adjusted $\beta^a$ | 95% CI | *p*-value |
| --- | --- | --- | --- | --- | --- | --- | --- |
| 1 | IL-1β, IL-8 | .2 | -.16; .57 | .27 | .07 | -.24; .38 | .67 |
| 2 | IL-4, IL-5, IL-6, IL-13 | -.23 | -.73; .28 | .37 | -.19 | -.72; .34 | .48 |
| 3 | IL-10, IL-12p70, IL-17A,<br>IL-23, IFN-γ, IL-2, GM-CSF | -.12 | -.69; .45 | .69 | .15 | -.43; .73 | .61 |

<sup>a</sup>: Adjusted for: maternal age, race/ethnicity, pre-pregnancy body mass index, gestational diabetes, parity, gestational age in weeks at blood draw, being in labor at the time of blood draw (yes / no), and time between blood draw & sample processing (hours).

## 4. Discussion

### 4.1 Principal findings

In the current study, we found that levels of individual cytokines and cytokine cluster Eigenvalues are not different between pregnant people with evidence of previous SARS-CoV-2 infection and those not recently infected.

### 4.2 Cytokine changes in SARS-CoV-2 infected pregnant people

Few earlier studies have focused on the immune response after SARS-CoV-2 infection during pregnancy. A recent study compared 90 pregnant people with a positive SARS-CoV-2 RT-PCR test of oropharyngeal or nasopharyngeal specimens with 90 pregnant people with a negative RT-PCR test, and showed that SARS-CoV-2 infection was associated with increased levels of IFN-γ in the third trimester, and decreased levels of IL-2, IL-10 and IL-17 in the first trimester and decreased levels of IL-2 in the second trimester [16]. Chen *et al*. showed that plasma cytokine levels of IL-2, IL-5, IL-10 and IL-12 in 11 pregnant people with acute SARS-CoV-2 infection were decreased compared to 10 healthy pregnant people [17]. Sherer *et al*. also compared SARS-CoV-2 infected pregnant people (n=33) and non-infected pregnant people (n=17) based on RT-PCR testing and did not find altered gene expression levels of *IL-1β* and *IL-6* [18]. However, they did find increased levels of *IL-1β* in maternal blood taken within 14 days of infection compared to samples taken 14 days after infection [18]. RT-PCR testing is commonly used to detect viral RNA of SARS-CoV-2 in respiratory secretions and a positive test is often considered an indicator of active viremia, although this is not guaranteed [39]. When measured in a person with a positive SARS-CoV-2 RT-PCR test, cytokine changes are likely a response to acute viral infection [40]. In the current study, we investigated whether SARS-CoV-2 infection during pregnancy based on anti-S IgG antibody detection is associated with persisting cytokine changes. We did not find changes in an extensive panel of cytokines compared between pregnant people with and without IgG antibodies to the SARS-CoV-2 spike protein, except for IL-23 at labor and delivery specifically. Together, these findings indicate that cytokine levels may be altered in pregnant people in the acute phase of SARS-CoV-2 infection, but are largely restored once the acute phase is resolved.

### 4.3 Consequences for pregnancy outcomes

Dysregulation of the immune system during pregnancy has been associated with adverse pregnancy outcomes [15, 41]. Since the start of the pandemic, studies have shown higher rates of preterm birth, preeclampsia and stillbirth, as well as increased maternal anxiety and mortality after SARS-CoV-2 infection during pregnancy [42]. It has been hypothesized that dysregulation of the immune system as a consequence of SARS-CoV-2 infection mediates the association between infection and pregnancy outcomes [43]. In the current study, we found no evidence for long-term maternal cytokine dysregulation in response to SARS-CoV-2 infection during pregnancy, except for decreased IL-23 levels in cases at labor and delivery. Although these results should be interpreted with care, as this was only a subgroup of our sample (n=75), this finding is of interest as IL-23 plays a key role in the development and differentiation of T helper 17 (TH17) cells through the activation of STAT3 [44], and dysregulation of the TH17/Treg balance is hypothesized to contribute to adverse pregnancy outcomes [43,45]. It should be noted that cytokines capture only part of the maternal immunology, and that other immune markers are important in understanding the effect of SARS-CoV-2 infection on the maternal immune system. Peripheral blood mononuclear cells (PBMC’s) are currently being collected from pregnant participants of the Generation C Study to enable a more extended analysis of the mediating role of the maternal immune system in SARS-CoV-2 exposed pregnant people, including T cell subsets.

### 4.4 Strengths and limitations

This study has several strengths. We used a serological assay with a high sensitivity (95%) and specificity (100%) to establish SARS-CoV-2 exposure [20]. This enabled unbiased recruitment of pregnant people with past SARS-CoV-2 infection, both with and without symptoms. Even though we do not know the exact timing of infection, based on time of sample collection and expected duration of antibodies we can infer that infection occurred during pregnancy. This study is the largest of its kind so far. We investigated a broad panel of cytokines, which allowed us to assess not only individual cytokine responses, but also cytokine clusters. The findings of the current study are also subject to some limitations. The difference in sample size between the second trimester samples compared to the third trimester and labor and delivery samples is due to the current study being an interim analysis of an ongoing prospective cohort study. Future studies based on the entire prospective cohort will include a larger sample for each timepoint. Pregnant people with false negative anti-S IgG antibodies may have been misclassified as unexposed since IgG production onset occurs one week after infection and even though studies have shown sustained IgG antibody levels after 30 weeks of infection [38], IgG antibody levels might incidentally revert to zero in the months after infection [25]. In addition, anti-S IgG antibodies were assessed only once in participants, at a median gestational age of 34 weeks (SD 38 days). The association between anti-S IgG antibody status and cytokine levels may differ throughout gestation. Repeated IgG measurements throughout gestation will provide more information on timing of infection and on the association between cytokines and changes in titer levels between assessments. Lastly, limited information was available regarding severity of infection. IgG antibody titer level was used as a proxy for disease severity as IgG antibody levels has been associated with disease severity [38]; however, this is not a one-to-one association as studies have shown cases of asymptomatic patients with high titers, and vice versa [46].

### 4.5 Conclusion

Our results from a case-control study comparing pregnant people with and without antibody evidence of SARS-CoV-2 infection found no indication of persisting maternal cytokine changes in response to SARS-CoV-2 infection during pregnancy. The current interim analysis suggests that the acute inflammatory response after SARS-CoV-2 infection during pregnancy may be restored to normal values within the pregnancy period. Results of the completed Generation C study from which these data are an interim analysis are expected end of 2022. Despite our finding that suggests that SARS-CoV-2 infection during pregnancy does not lead to lasting immune changes, more studies are needed to fully understand the risks of SARS-CoV-2 infection for pregnant and recently pregnant people. Prospective studies with repeated assessments throughout gestation, including groups with documented varying degrees of disease severity of SARS-CoV-2 infection, are needed to further validate these findings and understand the association between SARS-CoV-2 infection (severity) during pregnancy and maternal cytokine changes.

## Supporting information

Supplementary Table III

Supplementary Table VII

Supplementary Table VIII

## 5. Acknowledgements

We would like to thank all participants of the Generation C study for their cooperation and contribution to the research field. We would like to thank the Krammer Serology Core Study group, and especially Juan Manuel Carreño, Gagandeep Singh, Dominika Bielak and Daniel Stadlbauer for their help with running the SARS-CoV-2 ELISA’s. We would also like to thank the Mount Sinai Biorepository and Pathology Core, and especially Maryann Huie, Frances Avila, Ariane Benedetto, Anastasiya Dzhun, Shawn El Naggar, Bea Martin and Roni Sussman. We would like to thank several members of the U.S. Centers for Disease Control and Prevention (CDC) that have contributed to the interpretation of the data and have provided their feedback on the manuscript: Lauren B. Zapata, Van T. Tong, and Sascha R. Ellington.

