## Supplementary Figure 1 for "Maternal cytokine response after SARS-CoV-2 infection during pregnancy"

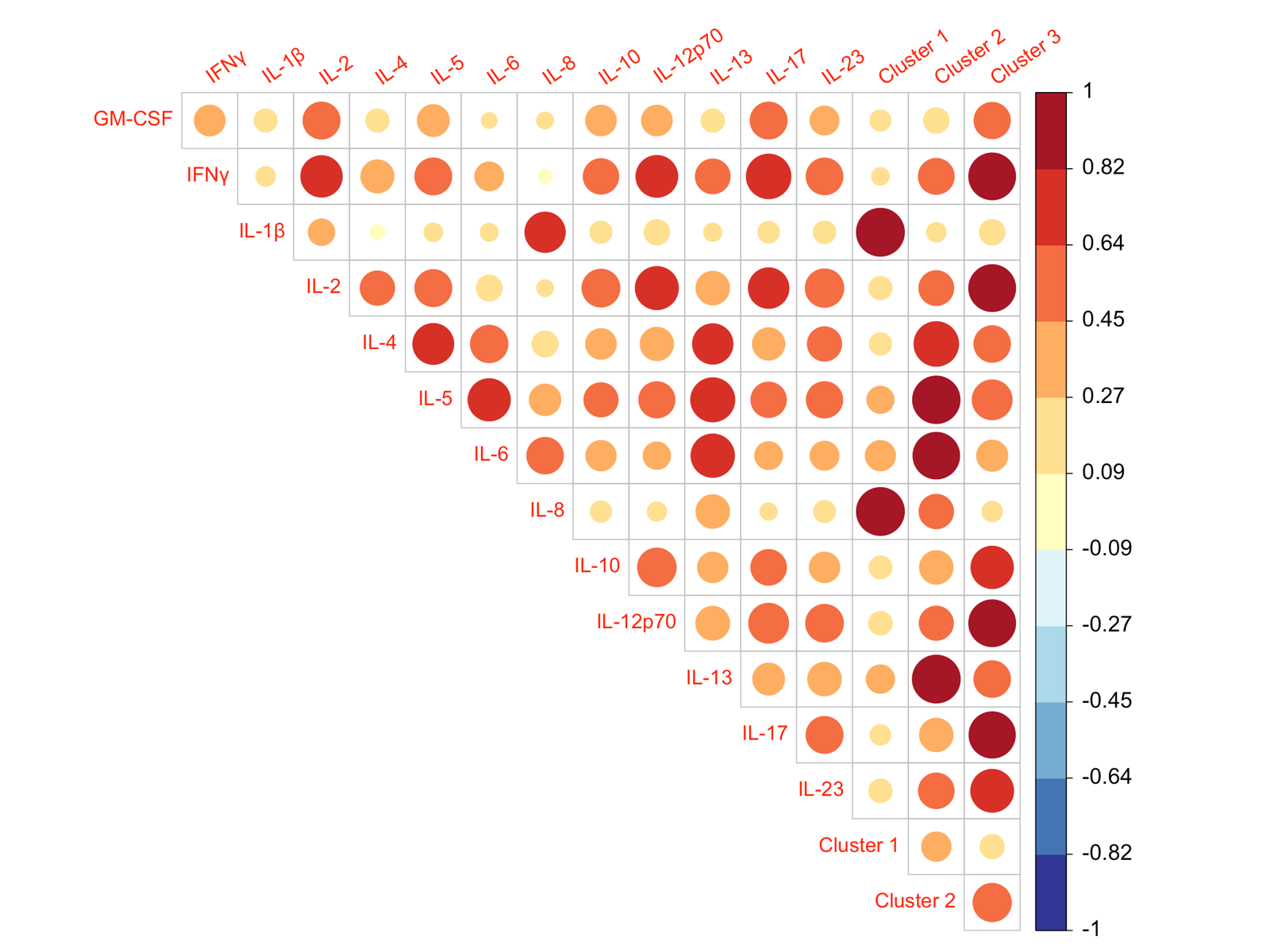

**Supplementary Figure 1.** Correlation plot of 13 cytokines assessed in the peripheral blood of cases (pregnant people with anti-S IgG antibodies, n=99) and controls (pregnant people with no anti-S IgG antibodies, n=99) and the cluster Eigenvalues. Darker colors indicate stronger positive correlations. The cytokines in cluster 1, 2 and 3 show the strongest correlation with their respective cluster Eigenvalues compared to the individual cytokines. GM-CSF = granulocyte-macrophage colony-stimulating factor, IFN-γ = Interferon, IL = interleukin.
