## Supplementary Figure 2 for "Maternal cytokine response after SARS-CoV-2 infection during pregnancy"

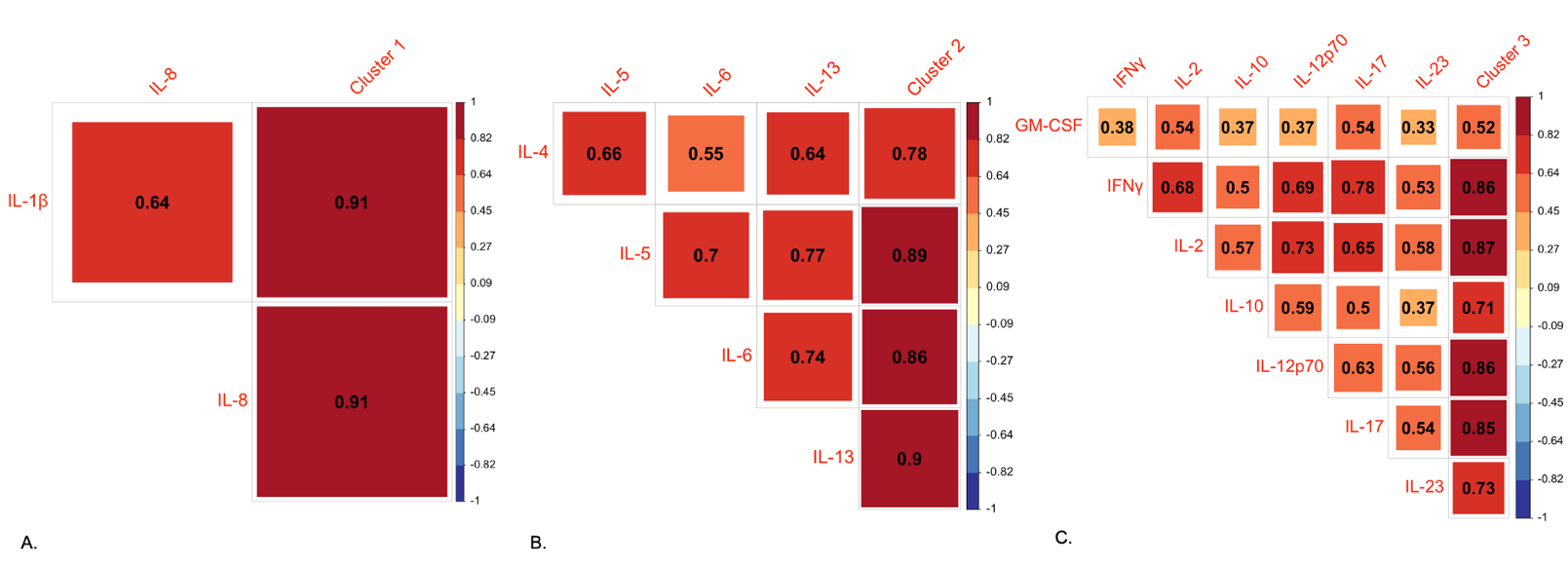

**Supplementary Figure 2.** Correlation plot of cluster 1 (A), cluster 2 (B) and cluster 3 (C) showing the concordance between the cytokines and their respective cluster Eigenvalues. Correlation coefficients rho are shown. Darker colors indicate stronger positive correlations. Cytokines in each cluster demonstrate the strongest level of association with their respective cluster Eigenvalues as compared to other cytokines, except for GM-CSF which shows strong correlation with IL-2 and IL-17. GM-CSF = granulocyte-macrophage colony-stimulating factor, IFN-γ = Interferon, IL = interleukin.
