## Supplementary Table I for "Maternal cytokine response after SARS-CoV-2 infection during pregnancy"

**Supplementary Table I. Descriptives of multiplex assay performance on 14 peripheral blood cytokines in blood samples of 200 pregnant people**

Descriptives of the multiplex assay performance of 14 peripheral blood cytokines, each measured in 200 samples. Two samples showed undetected cytokine values for IL-4 and IL-13 and were excluded from the analysis. All other samples (n=198) performed within the normal range. Cytokine values are measured in pg/ml. Pg = picogram, GM-CSF = granulocyte-macrophage colony-stimulating factor, IFN-γ = Interferon, IL = interleukin.

| Cytokine | N missing | Minimum (pg/ml) | Maximum (pg/ml) | Mean  (pg/ml) | Std. Deviation |
| --- | --- | --- | --- | --- | --- |
| GM-CSF | 0 | 12.6 | 1241.3 | 85.5 | 109.2 |
| IFN-γ | 0 | 4.3 | 70.1 | 14.9 | 8.1 |
| IL-1β | 0 | 0.3 | 71.9 | 4.3 | 8.1 |
| IL-2 | 0 | 1.4 | 25.2 | 4.9 | 2.5 |
| IL-4 | 1 | 2.9 | 258.8 | 24.0 | 28.8 |
| IL-5 | 0 | 0.4 | 11.1 | 2.0 | 1.5 |
| IL-6 | 0 | 0.2 | 28.0 | 2.2 | 3.9 |
| IL-8 | 0 | 1.5 | 1382.0 | 32.5 | 138.9 |
| IL-10 | 0 | 0.9 | 202.8 | 7.7 | 14.4 |
| IL-12p70 | 0 | 0.6 | 12.9 | 2.8 | 1.3 |
| IL-13 | 1 | 0.5 | 1694.7 | 16.4 | 120.2 |
| IL-17A | 0 | 2.0 | 36.1 | 8.7 | 4.7 |
| IL-23 | 0 | 73.1 | 3040.0 | 557.4 | 419.0 |
| TNFα | 0 | 1.0 | 19.5 | 4.1 | 2.1 |
