## Supplementary Table II for "Maternal cytokine response after SARS-CoV-2 infection during pregnancy"

**Supplementary Table II.** **Comparison of peripheral blood cytokine levels among pregnant people by anti-S IgG antibody titer levels**

Peripheral blood cytokine levels were compared among cases (pregnant people with anti-S IgG antibodies, n=99) divided by titer level into mild (1:80-1:400, n=37), moderate (1:800: 1:1600, n=46) and high (>1:1600, n=16), and anti-S IgG antibody negative pregnant people (n=99). Cytokine levels were not significantly different among cases with various titer levels, nor between cases and controls. GM-CSF = granulocyte-macrophage colony-stimulating factor, IFN-γ = Interferon, IL = interleukin.

| Cytokine | Anti-S IgG antibody negative  (n=99)^a^ | Anti-S IgG antibody positive  Mild titer  (n=37)^a^ | Anti-S IgG antibody positive  Moderate titer (n=46)^a^ | Anti-S IgG antibody positive  High titer  (n=16)^a^ | *p*-value^b^ |
| --- | --- | --- | --- | --- | --- |
| GM-CSF | 55.7 (54.8) | 56.98 (54.6) | 55.7 (56.9) | 55.3 (62.3) | .95 |
| IFN-γ | 12.9 (8.5) | 14.1 (8.8) | 12.3 (10.2) | 12.4 (5.5) | .68 |
| IL-1β | 1.7 (4.1) | 1.7 (5.4) | 1.9 (4.7) | 1.5 (.8) | .57 |
| IL-2 | 4.7 (2.6) | 4.3 (2.6) | 4.7 (3.0) | 4.5 (2.9) | .96 |
| IL-4 | 15.7 (10.3) | 17.2 (8.1) | 14.9 (12.0) | 15.2 (9.5) | .54 |
| IL-5 | 1.6 (.8) | 1.6 (.6) | 1.4 (.9) | 1.4 (1.2) | .58 |
| IL-6 | 0.9 (.7) | .93 (.95) | .87 (.61) | .86 (.56) | .72 |
| IL-8 | 6.7 (10.7) | 7.7 (9.1) | 7.1 (14.2) | 5.4 (5.5) | .52 |
| IL-10 | 6.2 (3.9) | 6.0 (3.0) | 5.99 (4.9) | 6.5 (4.3) | .73 |
| IL-12p70 | 2.5 (1.4) | 2.6 (1.7) | 2.3 (1.6) | 2.5 (1.0) | .91 |
| IL-13 | 3.9 (3.1) | 4.1 (5.3) | 3.8 (2.9) | 3.8 (2.2) | .76 |
| IL-17A | 7.1 (5.5) | 7.7 (5.3) | 6.6 (5.1) | 7.0 (5.6) | .54 |
| IL-23 | 402.7 (289.8) | 396.1 (301.3) | 415.9 (301.5) | 442.2 (301.2) | .40 |
| TNFα | 3.8 (2.6) | 3.3 (2.3) | 4.0 (2.5) | 3.9 (2.7) | .51 |

^a^: Medians and interquartile range values were reported for the non-normally distributed parameters.

^b^: Kruskall-Wallis test.
