## Supplementary Table III for "Maternal cytokine response after SARS-CoV-2 infection during pregnancy"

**Supplementary Table III. Comparison of peripheral blood cytokine levels between cases and controls in sensitivity analysis**

A sensitivity analysis was performed of n=125 blood samples of pregnant people in NYC, excluding samples obtained at labor and delivery (n=75). Peripheral blood cytokine levels were compared between cases (pregnant people with anti-S IgG antibodies, n=63) and controls (pregnant people with no anti-S IgG antibodies, n=62). Linear regression analyses were performed with plasma cytokine levels (pg/ml) as the dependent variables and SARS-CoV-2 status (anti-S IgG antibody positive / negative) as the exposure variable. Both unadjusted and adjusted beta coefficients are shown. Analyses were adjusted for covariates listed below. Similar to the analysis of the full sample (Table 2), cytokines were not significantly different between cases and controls for unadjusted nor adjusted linear regression analysis in the sensitivity analysis. GM-CSF = granulocyte-macrophage colony-stimulating factor, IFN-γ = Interferon, IL = interleukin.

| Cytokine | Unadjusted ß | 95% CI | *p*-value | Adjusted ß^a^ | 95% CI | *p*-value |
| --- | --- | --- | --- | --- | --- | --- |
| GM-CSF | -.02 | -.13; .09 | .71 | -.001 | -.11; .11 | .98 |
| IFN-γ | -.02 | -.09; .06 | .64 | -.004 | -.08; .07 | .92 |
| IL-1β | .07 | -.07; .2 | .32 | .07 | -.04; .18 | .21 |
| IL-2 | .004 | -.07; .08 | .92 | .03 | -.04; .11 | .37 |
| IL-4 | .04 | -.08; .15 | .5 | .06 | -.06; .18 | .32 |
| IL-5 | -.03 | -.11; .06 | .54 | -.02 | -.11; .08 | .72 |
| IL-6 | -.02 | -.17; .13 | .82 | -.02 | -.18; .13 | .79 |
| IL-8 | .01 | -.14; .15 | .92 | -.02 | -.14; .1 | .73 |
| IL-10 | .02 | -.06; .11 | .6 | .05 | -.04; .13 | .29 |
| IL-12p70 | .01 | -.06; .08 | .7 | .03 | -.04; .1 | .42 |
| IL-13 | -.06 | -.24; .12 | .51 | -.06 | -.25; .13 | .52 |
| IL-17A | -.01 | -.08; .06 | .8 | .01 | -.07; .08 | .82 |
| IL-23 | .02 | -.08; .11 | .74 | .03 | -.08; .13 | .62 |
| TNFα | -.02 | -.1; .06 | .62 | -.04 | -.11; .04 | .31 |

^a^: Adjusted for: maternal age, race/ethnicity, pre-pregnancy body mass index, gestational diabetes, parity, gestational age in weeks at blood draw, being in labor at the time of blood draw (yes / no), and time between blood draw & sample processing (hours).
