## Supplementary Table IV for "Maternal cytokine response after SARS-CoV-2 infection during pregnancy"

**Supplementary Table IV.** **Comparison of** **peripheral blood cytokine levels between cases and controls in the third trimester**

Peripheral blood cytokine levels were compared in the third trimester among cases (pregnant people with anti-S IgG antibodies, n=56) and controls (anti-S IgG antibody negative pregnant people, n=54). Linear regression analyses were performed with plasma cytokine levels (pg/ml) as the dependent variable and SARS-CoV-2 status (anti-S IgG antibody positive / negative) as the exposure variable. Analyses were adjusted for covariates listed below. Cytokine levels were not significantly different between cases and controls. GM-CSF = granulocyte-macrophage colony-stimulating factor, IFN-γ = Interferon, IL = interleukin.

| Cytokine | Adjusted ß^a^ | 95% CI | *p*-value |
| --- | --- | --- | --- |
| GM-CSF | .02 | -.1; .14 | .75 |
| IFN-γ | .001 | -.08; .08 | .98 |
| IL-1β | .04 | -.07; .14 | .52 |
| IL-2 | .05 | -.03; .13 | .21 |
| IL-4 | .08 | -.05; .2 | .24 |
| IL-5 | -.02 | -.12; .08 | .71 |
| IL-6 | -.02 | -.19; .15 | .81 |
| IL-8 | -.04 | -.16; .09 | .55 |
| IL-10 | .06 | -.04; .15 | .23 |
| IL-12p70 | .02 | -.05; .1 | .53 |
| IL-13 | -.06 | -.27; .15 | .57 |
| IL-17A | .02 | -.07; .1 | .7 |
| IL-23 | .03 | -.08; .14 | .56 |
| TNFα | -.03 | -.1; .04 | .34 |

^a^: Adjusted for: maternal age, race/ethnicity, pre-pregnancy body mass index, gestational diabetes, parity, gestational age in weeks at blood draw, being in labor at the time of blood draw (yes / no), and time between blood draw & sample processing (hours).
