## Supplementary Table V for "Maternal cytokine response after SARS-CoV-2 infection during pregnancy"

**Supplementary Table V.** **Comparison of** **peripheral blood cytokine levels between cases and controls at labor and delivery**

Peripheral blood cytokine levels were compared at labor and delivery among cases (pregnant people with anti-S IgG antibodies, n=37) and controls (anti-S IgG antibody negative pregnant people, n=38). Linear regression analyses were performed with plasma cytokine levels (pg/ml) as the dependent variable and SARS-CoV-2 status (anti-S IgG antibody positive / negative) as the exposure variable. Analyses were adjusted for covariates listed below. IL-23 levels were significantly lower in cases compared to controls at labor and delivery. Other cytokine levels were not significantly different between cases and controls. GM-CSF = granulocyte-macrophage colony-stimulating factor, IFN-γ = Interferon, IL = interleukin.

| Cytokine | Adjusted ß^a^ | 95% CI | *p*-value |
| --- | --- | --- | --- |
| GM-CSF | -.04 | -.2; .11 | .56 |
| IFN-γ | .02 | -.09; .13 | .72 |
| IL-1β | .07 | -.13; .28 | .47 |
| IL-2 | .02 | -.1; .13 | .75 |
| IL-4 | -.12 | -.29; .06 | .19 |
| IL-5 | -.04 | -.16; .08 | .49 |
| IL-6 | .06 | -.15; .26 | .59 |
| IL-8 | .08 | -.11; .28 | .41 |
| IL-10 | .01 | -.14; .17 | .88 |
| IL-12p70 | .004 | -.08; .09 | .93 |
| IL-13 | -.05 | -.25; .15 | .59 |
| IL-17A | .03 | -.09; .14 | .64 |
| IL-23 | -.15 | -.28; -.02 | *.03* |
| TNFα | .11 | .00; .22 | .06 |

^a^: Adjusted for: maternal age, race/ethnicity, pre-pregnancy body mass index, gestational diabetes, parity, gestational age in weeks at blood draw, being in labor at the time of blood draw (yes / no), and time between blood draw & sample processing (hours).
