## Supplementary Table VI for "Maternal cytokine response after SARS-CoV-2 infection during pregnancy"

**Supplementary Table VI.** **Correlation of cytokines measured in peripheral blood samples of 200 pregnant people**

Correlation matrix of 14 cytokines measured in peripheral blood samples of pregnant people in NYC (r = Spearman’s rho). TNFα shows a non-significant and mostly negative correlation with other cytokines (rho ranging between -0.288 and 0.39). GM-CSF = granulocyte-macrophage colony-stimulating factor, IFN-γ = Interferon, IL = interleukin.

| Cytokine |  | GM-CSF | IFNγ | IL-1β | IL-2 | IL-4 | IL-5 | IL-6 | IL-8 | IL-10 | IL-12p70 | IL-13 | IL-17A | IL-23 | TNFα |
| --- | --- | --- | --- | --- | --- | --- | --- | --- | --- | --- | --- | --- | --- | --- | --- |
| GM-CSF | r-value | 1 | .414** | .250** | .546** | .347** | .494** | .147* | .157* | .485** | .418** | .256** | .539** | .344** | -.132 |
|  | p-value | . | 0 | 0 | 0 | 0 | 0 | 0.037 | 0.027 | 0 | 0 | 0 | 0 | 0 | 0.063 |
|  | N | 200 | 200 | 200 | 200 | 199 | 200 | 200 | 200 | 200 | 200 | 199 | 200 | 200 | 200 |
| IFN-γ | r-value | .414** | 1 | .239** | .708** | .544** | .622** | .377** | .185** | .589** | .718** | .545** | .792** | .558** | -.194** |
|  | p-value | 0 | . | 0.001 | 0 | 0 | 0 | 0 | 0.009 | 0 | 0 | 0 | 0 | 0 | 0.006 |
|  | N | 200 | 200 | 200 | 200 | 199 | 200 | 200 | 200 | 200 | 200 | 199 | 200 | 200 | 200 |
| IL-1β | r-value | .250** | .239** | 1 | .415** | .261** | .269** | .233** | .574** | .303** | .384** | .231** | .250** | .318** | -.069 |
|  | p-value | 0 | 0.001 | . | 0 | 0 | 0 | 0.001 | 0 | 0 | 0 | 0.001 | 0 | 0 | 0.334 |
|  | N | 200 | 200 | 200 | 200 | 199 | 200 | 200 | 200 | 200 | 200 | 199 | 200 | 200 | 200 |
| IL-2 | r-value | .546** | .708** | .415** | 1 | .570** | .613** | .318** | .194** | .647** | .785** | .499** | .664** | .632** | -.300** |
|  | p-value | 0 | 0 | 0 | . | 0 | 0 | 0 | 0.006 | 0 | 0 | 0 | 0 | 0 | 0 |
|  | N | 200 | 200 | 200 | 200 | 199 | 200 | 200 | 200 | 200 | 200 | 199 | 200 | 200 | 200 |
| IL-4 | r-value | .347** | .544** | .261** | .570** | 1 | .683** | .464** | .352** | .490** | .549** | .676** | .509** | .460** | -.288** |
|  | p-value | 0 | 0 | 0 | 0 | . | 0 | 0 | 0 | 0 | 0 | 0 | 0 | 0 | 0 |
|  | N | 199 | 199 | 199 | 199 | 199 | 199 | 199 | 199 | 199 | 199 | 198 | 199 | 199 | 199 |
| IL-5 | r-value | .494** | .622** | .269** | .613** | .683** | 1 | .539** | .409** | .596** | .571** | .690** | .573** | .513** | -.222** |
|  | p-value | 0 | 0 | 0 | 0 | 0 | . | 0 | 0 | 0 | 0 | 0 | 0 | 0 | 0.002 |
|  | N | 200 | 200 | 200 | 200 | 199 | 200 | 200 | 200 | 200 | 200 | 199 | 200 | 200 | 200 |
| IL-6 | r-value | .147* | .377** | .233** | .318** | .464** | .539** | 1 | .516** | .381** | .372** | .601** | .301** | .305** | -.012 |
|  | p-value | 0.037 | 0 | 0.001 | 0 | 0 | 0 | . | 0 | 0 | 0 | 0 | 0 | 0 | 0.861 |
|  | N | 200 | 200 | 200 | 200 | 199 | 200 | 200 | 200 | 200 | 200 | 199 | 200 | 200 | 200 |
| IL-8 | r-value | .157* | .185** | .574** | .194** | .352** | .409** | .516** | 1 | .280** | .195** | .465** | .216** | .230** | .039 |
|  | p-value | 0.027 | 0.009 | 0 | 0.006 | 0 | 0 | 0 | . | 0 | 0.006 | 0 | 0.002 | 0.001 | 0.585 |
|  | N | 200 | 200 | 200 | 200 | 199 | 200 | 200 | 200 | 200 | 200 | 199 | 200 | 200 | 200 |
| IL-10 | r-value | .485** | .589** | .303** | .647** | .490** | .596** | .381** | .280** | 1 | .646** | .493** | .541** | .446** | -.200** |
|  | p-value | 0 | 0 | 0 | 0 | 0 | 0 | 0 | 0 | . | 0 | 0 | 0 | 0 | 0.004 |
|  | N | 200 | 200 | 200 | 200 | 199 | 200 | 200 | 200 | 200 | 200 | 199 | 200 | 200 | 200 |
| IL-12p70 | r-value | .418** | .718** | .384** | .785** | .549** | .571** | .372** | .195** | .646** | 1 | .508** | .640** | .591** | -.279** |
|  | p-value | 0 | 0 | 0 | 0 | 0 | 0 | 0 | 0.006 | 0 | . | 0 | 0 | 0 | 0 |
|  | N | 200 | 200 | 200 | 200 | 199 | 200 | 200 | 200 | 200 | 200 | 199 | 200 | 200 | 200 |
| IL-13 | r-value | .256** | .545** | .231** | .499** | .676** | .690** | .601** | .465** | .493** | .508** | 1 | .443** | .432** | -.161* |
|  | p-value | 0 | 0 | 0.001 | 0 | 0 | 0 | 0 | 0 | 0 | 0 | . | 0 | 0 | 0.023 |
|  | N | 199 | 199 | 199 | 199 | 198 | 199 | 199 | 199 | 199 | 199 | 199 | 199 | 199 | 199 |
| IL-17A | r-value | .539** | .792** | .250** | .664** | .509** | .573** | .301** | .216** | .541** | .640** | .443** | 1 | .565** | -.186** |
|  | p-value | 0 | 0 | 0 | 0 | 0 | 0 | 0 | 0.002 | 0 | 0 | 0 | . | 0 | 0.008 |
|  | N | 200 | 200 | 200 | 200 | 199 | 200 | 200 | 200 | 200 | 200 | 199 | 200 | 200 | 200 |
| IL-23 | r-value | .344** | .558** | .318** | .632** | .460** | .513** | .305** | .230** | .446** | .591** | .432** | .565** | 1 | -.223** |
|  | p-value | 0 | 0 | 0 | 0 | 0 | 0 | 0 | 0.001 | 0 | 0 | 0 | 0 | . | 0.002 |
|  | N | 200 | 200 | 200 | 200 | 199 | 200 | 200 | 200 | 200 | 200 | 199 | 200 | 200 | 200 |
| TNFα | r-value | -0.132 | -.194** | -0.069 | -.300** | -.288** | -.222** | -0.012 | 0.039 | -.200** | -.279** | -.161* | -.186** | -.223** | 1 |
|  | p-value | 0.063 | 0.006 | 0.334 | 0 | 0 | 0.002 | 0.861 | 0.585 | 0.004 | 0 | 0.023 | 0.008 | 0.002 | . |
|  | N | 200 | 200 | 200 | 200 | 199 | 200 | 200 | 200 | 200 | 200 | 199 | 200 | 200 | 200 |
