## Supplementary Table VII for "Maternal cytokine response after SARS-CoV-2 infection during pregnancy"

**Supplementary Table VII.** **Pattern matrix of Principal Component Analysis of 13 cytokines** **measured in peripheral blood samples of 200 pregnant people**

A principal component analysis was performed on 13 cytokines measured in the peripheral blood samples of 200 pregnant people. The Oblimin with Kaiser Normalization was used. The rotation converged in 6 iterations. Component 1, 2 and 3 represent cluster 1, 2 and 3 respectively. IL-1β and IL-8 showed highest loading on component 1; IL-4, IL-5, IL-6 and IL-13 formed component 2; IL-10, IL17A, IFN-γ, IL-2, IL-12p70, IL-23 and GM-CSF showed highest loading on component 3. GM-CSF = granulocyte-macrophage colony-stimulating factor, IFN-γ = Interferon, IL = interleukin.

| Cytokine | Component | | |
| --- | --- | --- | --- |
|  | 1 | 2 | 3 |
| IL-17A | -0.033 | -0.023 | 0.861 |
| IFN-γ | -0.04 | -0.044 | 0.846 |
| IL-2 | 0.058 | -0.021 | 0.766 |
| IL-12p70 | 0.044 | -0.101 | 0.697 |
| IL-10 | -0.022 | -0.094 | 0.659 |
| GM-CSF | 0.043 | 0.217 | 0.555 |
| IL-23 | 0.041 | -0.254 | 0.553 |
| IL-13 | -0.046 | -0.8 | 0.226 |
| IL-6 | 0.044 | -0.915 | -0.098 |
| IL-5 | 0.014 | -0.849 | 0.155 |
| IL-4 | -0.027 | -0.784 | -0.029 |
| IL-8 | 0.858 | -0.114 | -0.092 |
| IL-1β | 0.857 | 0.051 | 0.088 |
