## Supplementary Table VIII for "Maternal cytokine response after SARS-CoV-2 infection during pregnancy"

**Supplementary Table VIII. Comparison of peripheral blood cytokine clusters among pregnant people by anti-S IgG antibody titer levels**

The Eigenvalues of three cytokine clusters were compared among cases (pregnant people with anti-S IgG antibodies, n=99) divided by titer level into mild (1:80-1:400, n=37), moderate (1:800: 1:1600, n=46) and high (>1:1600, n=16), and anti-S IgG antibody negative pregnant people (n=99). The Eigenvalues of the clusters were not significantly different among cases with various titer levels, nor between cases and controls. GM-CSF = granulocyte-macrophage colony-stimulating factor, IFN-γ = Interferon, IL = interleukin.

| Cluster | Cytokines in cluster | Anti-S IgG antibody negative  (n=99)^a^ | Anti-S IgG antibody positive  Mild titer  (n=37)^a^ | Anti-S IgG antibody positive  Moderate titer (n=46)^a^ | Anti-S IgG antibody positive  High titer  (n=16)^a^ | *p*-value^b^ |
| --- | --- | --- | --- | --- | --- | --- |
| 1 | IL-1β, IL-8 | -.1 (1.2) | .11 (1.2) | .16 (1.4) | -.2 (1.5) | .51 |
| 2 | IL-4, IL-5, IL-6, IL-13 | .11 (1.8) | .13 (1.6) | -.25 (1.8) | -.29 (1.4) | .48 |
| 3 | IL-10, IL-12p70, IL-17A, IL-23, IFN-γ, IL-2, GM-CSF | .11 (2.0) | .19 (1.9) | -.31 (2.1) | .01 (1.7) | .76 |

^a^: Mean and standard deviation values were reported for the non-normally distributed parameters.

^b^: Kruskall-Wallis test.
